# *tadar*: an R/Bioconductor package to reduce eQTL noise in differential expression analysis

**DOI:** 10.1101/2025.02.11.637777

**Authors:** Lachlan Baer, Michael Lardelli, Stevie M. Pederson

**Affiliations:** School of Biological Sciences, University of Adelaide, Adelaide, South Australia, Australia; Black Ochre Data Labs, Indigenous Genomics, The Kids Research Institute Australia, Adelaide, South Australia, Australia; School of Biomedicine, University of Adelaide, Adelaide, South Australia, Australia; John Curtin School of Medical Research, Australian National University, Canberra, Australian Capital Territory, Australia

**Keywords:** RNA-Sequencing, eQTL, differential expression, bias, DAR, tadar

## Abstract

Despite the relative maturity of bulk RNA-sequencing, compared to more recent developments such as single-cell and spatial RNA-sequencing, biases which impact data analysis continue to surface. One such bias, termed “Differential Allelic Representation” (DAR), is particularly evident when experimental samples are taken from non-isogenic genetic backgrounds. DAR is an uneven distribution of polymorphic loci between groups of experimental samples undergoing differential gene expression (DGE) analysis. When unequally represented polymorphic loci are also expression quantitative trait loci (eQTLs), DAR can lead to differences in gene expression which are not directly relevant to the primary research objectives. To mitigate DAR in both new and existing datsets, we introduce *tadar*, a Bioconductor package designed to facilitate transcriptome analysis by accounting for differential allelic representation. *tadar* implements a methodology that calculates a DAR metric at each polymorphic locus across the genome, which then serves as a predictive measure of a locus’ potential to cause eQTL-driven expression differences. This metric is then used to reduce eQTL noise in bulk RNA-Sequencing data.

## Introduction

Expression Quantitative Trait Loci (eQTLs) are regions of the genome where DNA sequence variation impacts the expression of one or more genes (Brem et al., 2002; Schadt et al., 2003), and have been extensively studied due to their relevance in determining differential disease risk among individuals (Pritchard, 2002). DNA variants are commonly examined using Genome-Wide Association Studies (GWAS), where variant loci are identified that show statistically significant associations with a phenotype, such as disease (Hardy and Singleton, 2009; Hirschhorn et al., 2002). Such studies have revealed that the majority of variants lie within non-coding regions, suggesting an involvement in gene regulation (Manolio et al., 2009; Maurano et al., 2012). eQTL mapping employs a similar approach to GWAS in order to identify variant loci which impact the expression of specific genes (Westra and Franke, 2014). eQTLs are classified as cis (locally acting) or trans (distantly acting). Cis-eQTLs are conventionally found within 1 megabase (Mb) of a transcription start site (TSS), while trans-eQTLs are primarily located at least 5 Mb up- or downstream of a TSS (Nica and Dermitzakis, 2013).

eQTLs can pose a challenge for differential gene expression (DGE) analysis, when there is an unequal representation of eQTL loci between experimental sample groups being studied. This situation results from localised genetic diversity between the groups, and we have previously defined this as Differential Allelic Representation (DAR) (Baer et al., 2024). Commonly, when using animal models to investigate the impact of a gene mutation, the progeny of a single mating (or as few matings as possible) constitute the biological sample groups for investigation, with the intent of minimising genetic variability. Ideally, the mating involves a completely isogenic parental pair of animals, as is commonly seen in mouse experiments which use highly inbred strains (Simon et al., 2013). However, when the use of isogenic parental animals is not feasible, any genetic differences between parents are passed on to their progeny through Mendelian inheritance, with shuffling of haplotype blocks due to recombination. When segregated into experimental sample groups for comparison, such as for DGE analysis, the impacts of DAR can then become a confounding factor (Baer et al., 2024). In treatment vs. control experiments, the influence of DAR on DGE is a result of parental genomic diversity and the stochasticity of the above mechanisms during meiosis. However, in studies that require definition of experimental groups by the presence or absence of a genetic feature, such as an induced mutation in a specific gene, the selection for this feature can drive reduced genetic diversity in regions proximal to the selected locus within each group, thereby exacerbating DAR between the two groups in this region. When eQTLs lie within regions of high DAR, expression differences can be observed for genes regulated by these eQTLs. Whilst these observations are indeed indicative of differential expression, they are more correctly considered to be type I errors, given the intended context of the null hypothesis is that there is no differential expression between the two sample groups due to the specific genetic feature being studied.

DAR analysis is the process of calculating and assigning a metric at each single nucleotide polymorphism (SNP) across a genome reflecting the bias in its representation between two sample groups. DAR analysis requires only variant calls as input, allowing for its utilisation in experiments where only RNA-Sequencing (RNA-Seq) data is available, but also those with complementary DNA-sequencing (DNA-seq) data. Where DNA-seq data is available, this is the preferable choice for the generation of variant calls as it results in a more extensive representation of DAR across non-coding regions which may harbour eQTLs, and is less impacted by allelic bias. Where only RNA-Seq data is available, genetic variation information is indeed lost from a large portion of the genome. However, due to genetic linkage, loci close together on the same chromosome, i.e. in linkage disequilibrium, have a greater than fifty percent chance of being co-inherited from the same parental haplotype (Slatkin, 2008). Therefore, although DAR is calculated for a single-base locus, DAR estimates obtained from RNA-Seq are still able to partially bridge regions of missing information, such as non-transcribed regions of the genome. This understanding, along with the knowledge that cis-eQTLs tend to act in relatively close proximity to the genes they regulate, allows for the effective identification and evaluation of genes which are likely to be impacted by cis-eQTL effects.

In experiments where DAR is a likely confounder, DAR analysis can be employed to determine regions of the genome prone to eQTL-driven false positives. At a minimum, DAR analysis provides a basis for scepticism regarding the experimental relevance of differentially expressed genes or transcripts that lie within high DAR regions. However, we also propose and implement a method to directly moderate the p-values returned by differential expression testing. This approach can reduce the statistical significance of potentially problematic features, whilst ensuring no false positives are introduced.

Here we present *tadar*, a *Bioconductor* (Huber et al., 2015) package to account for the impacts of DAR during transcriptomic analysis. At its core, DAR analysis utilises the Euclidean distance metric to measure the extent of genetic variation between two groups of samples. In the following sections we outline the full methodology of DAR analysis and its implementation using *tadar*.

## Methods

### Package overview

The *tadar* package provides key functions required to perform DAR analysis, with each function performing a discrete step of the sequential process. This keeps less of the process hidden from view, intentionally exposing users to intermediate objects, allowing increased transparency, improved decision making and reinforcing understanding, instead of simply offering a black-box approach. The package has been designed for ease-of-use, requiring only a few lines of code to perform a complete analysis. The functions are also compatible with the base *R* pipe operator (R Core Team, 2024). An example of the code required to perform a complete DAR analysis with *tadar* can be seen below:

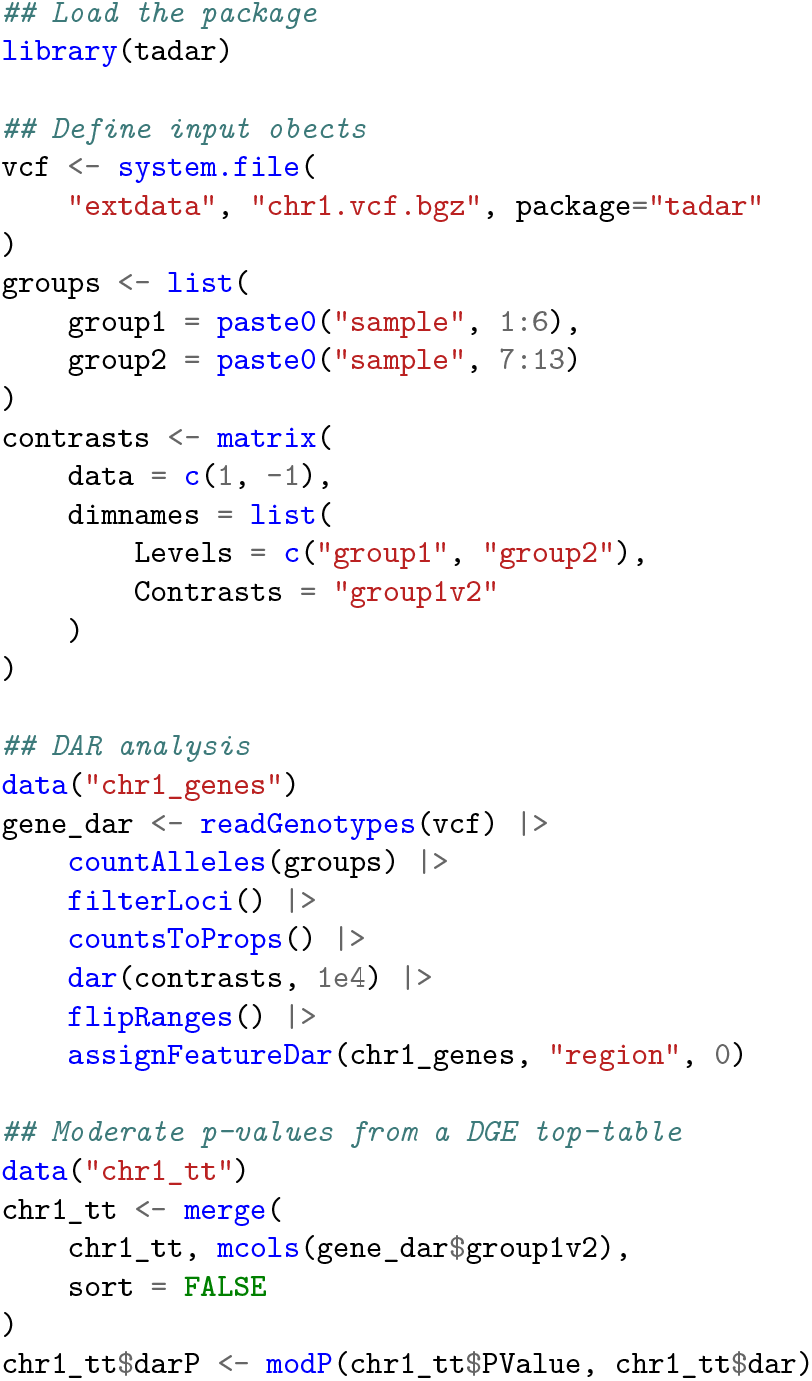

A detailed explanation of all functions used for the individual steps within the DAR analysis section is given below. These are also described in the *tadar* vignette hosted on the Bioconductor website. We first provide a path to our VCF, then define which samples represented in the VCF belong to a particular experimental group using a named list, and with each element simply being a vector of sample names. The specific contrasts can also be defined during the initial setup.

### Importing VCF data into the R environment

*tadar* relies on the well-established GRanges and GRangesList classes of the GenomicRanges package (Lawrence et al., 2013) throughout its workflow. The *readGenotypes* function is a wrapper for the *readVCF* function from the VariantAnnotation package (Obenchain et al., 2014), designed to import VCF data into the R environment. *readGenotypes* takes a multi-sample VCF file path as input and, by default, selectively parses only the CHROM, POS, ID, REF, and GT fields, as the minimum required data for DAR analysis. Since VCF files can be very large, this selective parsing significantly speeds up computational time compared to parsing the entire VCF contents. The resulting GRanges object contains the ranges corresponding to genomic coordinates where variants have been called in at least one sample. Genotype information is stored in the metadata columns, with each column representing a single sample.

### Summarising variation within sample groups

In order to capture allelic diversity within each group, *countAlleles* takes the initial parsed genotypes and summarises allele counts within each sample group, returning a GRangesList with length corresponding to the number of experimental groups. The metadata columns of each GRanges element include the counts of A, C, G, and T nucleotides at each locus, using *n_0* to denote the count of reference alleles, with *n_1* to *n_3* allowing for all four nucleotides to be represented within a group. However, given the nature of most parental crosses, only two alleles are typically observed at a locus. The function also summarises the number of called and missing genotypes, which are able to be used for filtering of loci, as described in the following section.

### Filtering loci and Obtaining Proportions

Underrepresented loci should be removed from the dataset as they may bias downstream calculation of the DAR metric. The default settings used in the *tadar* package are to retain any loci within a group where the number of samples with called genotypes exceeds 50%, however, users can specify their own filtering criteria when using *filterLoci*. The filter argument supports quasiquotation (Henry and Wickham, 2024), allowing users to define filtering criteria using any of the column names contained in the GRanges metadata. As a result of filtering, the number of represented loci in the GRangesList elements may vary between sample groups. Given that the number of samples within a sample group may differ, and the number of called genotypes may also vary from locus to locus within a group, count columns (*n_i* ) are then converted to proportions (*prop_i*), using *countsToProps*, to ensure allele counts are comparable across sample groups.

### Calculating DAR

The DAR metric (*d*) is defined as the Euclidean distance between the two vectors containing the allelic proportions at a given locus, divided by 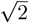 to ensure the value always lies on [0, 1], for easy interpretation. For a given locus and sample group comparison (e.g. *x* vs. *y* ), this is summarised with the equation,

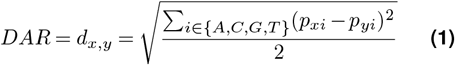

where *d*_*x*,*y*_ represents the DAR value between sample groups *x* and *y* at a single locus. *p*_*xi*_ and *p*_*yi*_ represent the proportions of each allele (A, C, G or T) observed at the locus in sample groups *x* and *y* respectively. *tadar* encapsulates this methodology with the *dar* function, taking a GRangesList as input, with each list element containing the proportions, and with the contrasts as specified.

The *dar* function reports DAR values for the intersection of loci represented in both groups, internally ignoring any loci exclusive to a single group. The number of retained loci reflects the stringency of filtering applied in the preceding step. A strict filter results in fewer loci but provides an overall increase in accuracy compared to a permissive filter that retains more samples with missing genotype calls.

In addition to calculating the metric at a given locus, the user can specify either of the “region_fixed” or “region_loci” arguments to produce a smoothed version of the metric. Passing a window size to “region_fixed” establishes a window of fixed size around each locus, with values averaged within each window. As an alternative, passing an integer to “region_loci” establishes an elastic region to average the specified number of loci, such that this number of values surrounding each origin locus are included. It should be noted that the latter choice likely results in genomic distances either side of the origin locus which are not symmetric. Both methods offer specific advantages and the choice is left to the user’s discretion and can be informed by the features of a given data set and for each individual application.

### Switching between origin and region representations of DAR

The output of the *dar* function reports two values for each locus, labelled as *dar_origin* and *dar_region* in the metadata columns of the resulting GRanges object. Raw, locus-specific DAR values are provided in the *dar_origin* column, whilst the *dar_region* values are the smoothed DAR values described above. The initial output of the *dar* function contains single-nucleotide ranges associated with the *dar_origin* values, and to “flip” these ranges to be the complete ranges used during smoothing, the *flipRanges* function is provided. This function allows reversible switching between the range representations, so users can change between the *dar_origin* loci or the smoothed region.

### Assigning DAR values to features

To interpret how features are impacted to eQTL-influenced differential expression, DAR values need to be assigned to the features (e.g. genes) being tested for differential expression using the *assignFeatureDar* function. This function requires two GRanges objects: one from the *dar* function output and another specifying feature annotations through the *features* argument, allowing application of DAR analysis to any type of genomic feature.

There are two options for assigning DAR values. First, the user can assign a DAR value to features containing an origin DAR value exclusively within a feature’s range. This method requires no modifications to the GRanges object prior to assignment. Alternatively, if the user prefers to assign values from a region surrounding the feature, they must first “flip” the ranges using the *flipRanges* function. This changes the ranges from those representing origin values to those representing smoothed regions, typically resulting in more features being assigned DAR values.

For each feature, *assignFeatureDar* calculates the mean of origin DAR values with positions that overlap each feature’s range. When origin ranges are used, the assigned DAR values represent the average DAR confined specifically to the feature. This approach is limited by its reliance on the detection of a variant within the feature itself, potentially missing eQTLs located outside the feature’s range. Alternatively, if the ranges are flipped to represent regions, this constructs a region surrounding the feature that is the same size as specified for smoothing DAR values. In this use case, origin DAR values are used for assignment to avoid additional averaging of an already averaged value.

### DAR moderation of p-values

As the final step of DAR analysis, we propose a method that leverages assigned DAR values to directly adjust feature level *p*-values from differential expression testing. The goal of this approach is to minimise noise introduced by eQTL-driven artefacts of differential expression, reducing the number of false positives without the introduction of further Type I errors.

Under the null hypothesis (*H*_0_), which assumes no differential expression between the two groups, *p*-values follow a uniform (𝒰) distribution on (0, 1), which is a special case of the Beta distribution, governed by the beta function:

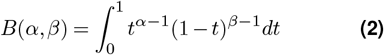

where *α* > 0 and *β* > 0 are parameters governing the shape of the distribution. When both *α* and *β* and are set to 1, the distribution is 𝒰 (0, 1).

In the presence of DAR, we suggest that under the most relevant specification of *H*_0_, i.e. the induced genetic change has no impact on gene expression, *p*-values are no longer uniformly distributed but are instead right-skewed, due to unequally distributed eQTLs. Additionally, this distribution will become more right-skewed as DAR approaches 1. These suggestions were supported by our observations across five different datasets where DAR was a confounder to differential expression (see Supplementary Information). By fitting both parameters of the Beta distribution as a function of DAR, we observed an approximately linear and decreasing relationship between DAR and the *α* parameter, whilst the *β* parameter was consistently estimated around the value 1. Thus, by modelling the *α* parameter of the beta distribution as a linear function of DAR, whilst holding *β* = 1, we can approximate the distribution from which *p*-values are drawn in a locus- or feature-specific manner. Feature-specific p-values can then be moderated using the appropriate DAR-based Beta distribution, which will lead to more conservative p-values in the presence of high DAR, whilst for features not associated with any notable DAR, p-values will remain unchanged. In this way, features which appear differentially expressed in a comparison, but where differential expression is driven by DAR-associated eQTLs rather than the intended genetic feature, can be reduced in their apparent significance.

This approach is implemented in the *modP* function of the *tadar* package. The *slope* argument has a default value of *a* = −1.8, chosen to be stringent. We recommend values −1 *< a <* −2, however users can follow the procedures outlined in the Supplementary Information to optimise this parameter for their own data. Additionally, the value *α* is only modified when the DAR value (*d*) is above a given threshold *b*, setting *b* = 0.1 by default. Thus for each feature *i* with an associated DAR value *d*_*i*_, the *α* parameter of the beta distribution is then modelled as the piece-wise linear function:

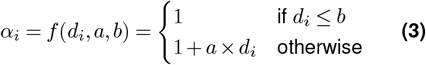

These fitted values for *α*_*i*_ are then used to obtain the probability of observing a given *p*-value (*p*_*i*_) under the distribution *B*(*α*_*i*_, 1), thereby moderating the set of *p*-values from the relevant DGE analysis. This process leads to an overall decrease in significance across the dataset, with genes located in high DAR regions being most affected, and those in low DAR regions being mostly unchanged. Importantly, features will only ever lose significance, ensuring that the moderation does not introduce further false positives. However, the ranking of each feature is likely to better reflect the intended biological question.

As a result, the moderated data becomes more suitable for enrichment techniques, such as Gene Set Enrichment Analysis (GSEA). By reducing the prominence of false positives, these features are pushed further towards the middle of the ranked list, which is commonly constructed based on the ordering of *p*-values and the direction of effect sizes. This improves the reliability of enrichment analyses by focusing attention on biologically relevant genes.

## Discussion

The package *tadar* provides a user-friendly interface that makes standardised DAR analysis accessible to both experienced bioinformaticians and researchers with only a basic understanding of R programming. The package requires a multi-sample VCF file containing SNP calls as input, for which we recommend following the GATK Best Practices Workflows (van der Auwera and O’Connor, 2020). A VCF file can be generated from either DNA-Seq or RNA-Seq data. DNA-Seq data offers a comprehensive view of DAR across the entire genome, enabling calculations of DAR in genomic regions that would otherwise be missed with only RNA-Seq data. This is particularly relevant since, as disease studies have shown, eQTLs often reside in non-coding regions (Manolio et al., 2009; Maurano et al., 2012). However, variant calls can also be derived from RNA-Seq data. Viable estimates of DAR can be obtained from RNA-Seq data by considering the influences on DAR due to genetic linkage and while working within a region-based context. Specifically, the DAR calculated from genes within a region can serve as a useful approximation for the DAR in nearby regions. For example, White et al. (White et al., 2022) determined the haplotypes of 96 SAT line zebrafish RNA-seq samples in 1 Mbp windows. This resulted in 82 regions across the genome where contiguous haplotypes could be resolved to the original strain of an intercross between one double haploid AB fish and one double haploid Tübingen fish, irrespective of the multiple generations of incrossing that had occurred. This principle underlies the motivation for calculating smoothed DAR values. As a result, DAR analysis remains applicable to existing RNA-Seq datasets, even in the absence of complementary DNA-Seq data.

To further streamline and standardise the process before DAR analysis, we also provide a workflow based on the GATK workflow for RNA-seq short variant discovery (SNPs + Indels), ensuring ease of use and reproducibility for users. This is accessible from the Snakemake (Mölder et al., 2021) workflow catalog or the GitHub repository https://github.com/baerlachlan/smk-rnaseq-gatk-variants. Whilst DAR analysis effectively addresses the confounding effects of cis-eQTL representation on differential expression outcomes, certain limitations remain. If a trans-eQTL is present in the data and is biasing gene expression within a region far from the variant site, DAR analysis is not able to appropriately account for this influence. However, given that trans-eQTLs typically exhibit smaller effect sizes (Pierce et al., 2014; Grundberg et al., 2012), this is considered to be a minor limitation.

Another potential limitation is the possibility of identifying true positives for differential expression within regions of high DAR, as may be likely for an induced mutation which directly impacts expression of the mutated gene. In such cases, the moderation of *p*-values can result in these true positives being assigned more conservative *p*-values. However, genes with substantial effect sizes that are truly differentially expressed are unlikely to lose statistical significance due to this process, and the un-moderated p-values are still able to provide important insights. The overall ranking of genes by differential expression following the moderation should be improved after accounting for DAR, as it better reflects the biological relevance of the results. Importantly, this refined ranking enhances the utility of the data in downstream analyses, such as Gene Set Enrichment Analysis (GSEA). Since GSEA relies on significance ranking and direction of effect sizes rather than *p*-values, the modifications from DAR analysis should lead to more accurate and interpretable outcomes for enrichment studies.

Our proposed methodology serves as a foundation for improving the interpretation of differential expression analysis in the presence of DAR. There are several future directions that could further enhance these interpretations, including the potential application of Bayesian approaches for modelling the values used in *B*(*α*_*i*_, *β*), and may provide a more robust framework for addressing the issue. This approach may also be applicable to other potential and identifiable sources of false positive results within DGE analyses.

*tadar*, the software implementing the described methodology, is available as an open-source package via the Bioconductor repository.

## Acknowledgements

The authors wish to thank Dr Jono Tuke for discussions regarding statistical methodology.

## Bibliography

Baer, L., Barthelson, K., Postlethwait, J. H., Adelson, D. L., Pederson, S. M., and Lardelli, M. (2024). Differential allelic representation (DAR) identifies candidate eQTLs and improves transcriptome analysis. PLOS Computational Biology, 20(2):e1011868. doi: 10.1371/journal.pcbi.1011868.

Brem, R. B., Yvert, G., Clinton, R., and Kruglyak, L. (2002). Genetic dissection of transcriptional regulation in budding yeast. Science, 296(5568):752–755. doi: 10.1126/science.1069516.

Grundberg, E., Small, K. S., Hedman, Å.K., Nica, A. C., Buil, A., Keildson, S., Bell, J. T., Yang, T.-P., Meduri, E., Barrett, A., Nisbett, J., Sekowska, M., Wilk, A., Shin, S.-Y., Glass, D., Travers, M., Min, J. L., Ring, S., Ho, K., Thorleifsson, G., Kong, A., Thorsteindottir, U., Ainali, C., Dimas, A. S., Hassanali, N., Ingle, C., Knowles, D., Krestyaninova, M., Lowe, C. E., Di Meglio, P., Montgomery, S. B., Parts, L., Potter, S., Surdulescu, G., Tsaprouni, L., Tsoka, S., Bataille, V., Durbin, R., Nestle, F. O., O’Rahilly, S., Soranzo, N., Lindgren, C. M., Zondervan, K. T., Ahmadi, K. R., Schadt, E. E., Stefansson, K., Smith, G. D., McCarthy, M. I., Deloukas, P., Dermitzakis, E. T., Spector, T. D., and The Multiple Tissue Human Expression Resource (MuTHER) Consortium. (2012). Mapping cis- and transregulatory effects across multiple tissues in twins. Nature Genetics, 44(10):1084–1089. doi: 10.1038/ng.2394.

Hardy, J. and Singleton, A. (2009). Genomewide Association Studies and Human Disease. New England Journal of Medicine, 360(17):1759–1768. doi: 10.1056/NEJMra0808700.

Henry, L. and Wickham, H. rlang: Functions for Base Types and Core R and ‘Tidyverse’ Features, (2024). URL https://CRAN.R-project.org/package=rlang. R package version 1.1.4.

Hirschhorn, J. N., Lohmueller, K., Byrne, E., and Hirschhorn, K. (2002). A comprehensive review of genetic association studies. Genetics in Medicine, 4(2):45–61. doi: 10.1097/00125817-200203000-00002.

Huber, W., Carey, V. J., Gentleman, R., Anders, S., Carlson, M., Carvalho, B. S., Bravo, H. C., Davis, S., Gatto, L., Girke, T., Gottardo, R., Hahne, F., Hansen, K. D., Irizarry, R. A., Lawrence, M., Love, M. I., MacDonald, J., Obenchain, V., Ole’s, A. K., Pag’es, H., Reyes, A., Shannon, P., Smyth, G. K., Tenenbaum, D., Waldron, L., and Morgan, M. (2015). Orchestrating high-throughput genomic analysis with Bioconductor. Nature Methods, 12 (2):115–121. doi: 10.1038/nmeth.3252.

Lawrence, M., Huber, W., Pagès, H., Aboyoun, P., Carlson, M., Gentleman, R., Morgan, M. T., and Carey, V. J. (2013). Software for Computing and Annotating Genomic Ranges. PLoS Computational Biology, 9(8):e1003118. doi: 10.1371/journal.pcbi.1003118.

Manolio, T. A., Collins, F. S., Cox, N. J., Goldstein, D. B., Hindorff, L. A., Hunter, D. J., McCarthy, M. I., Ramos, E. M., Cardon, L. R., Chakravarti, A., Cho, J. H., Guttmacher, A. E., Kong, A., Kruglyak, L., Mardis, E., Rotimi, C. N., Slatkin, M., Valle, D., Whittemore, A. S., Boehnke, M., Clark, A. G., Eichler, E. E., Gibson, G., Haines, J. L., Mackay, T. F. C., McCarroll, S. A., and Visscher, P. M. (2009). Finding the missing heritability of complex diseases. Nature, 461(7265):747–753. doi: 10.1038/nature08494.

Maurano, M. T., Humbert, R., Rynes, E., Thurman, R. E., Haugen, E., Wang, H., Reynolds, A. P., Sandstrom, R., Qu, H., Brody, J., Shafer, A., Neri, F., Lee, K., Kutyavin, T., Stehling-Sun, S., Johnson, A. K., Canfield, T. K., Giste, E., Diegel, M., Bates, D., Hansen, R. S., Neph, S., Sabo, P. J., Heimfeld, S., Raubitschek, A., Ziegler, S., Cotsapas, C., Sotoodehnia, N., Glass, I., Sunyaev, S. R., Kaul, R., and Stamatoyannopoulos, J. A. (2012). Systematic Localization of Common Disease-Associated Variation in Regulatory DNA. Science, 337(6099):1190–1195. doi: 10.1126/science.1222794.

Mölder, F., Jablonski, K. P., Letcher, B., Hall, M. B., Tomkins-Tinch, C. H., Sochat, V., Forster, J., Lee, S., Twardziok, S. O., Kanitz, A., Wilm, A., Holtgrewe, M., Rahmann, S., Nahnsen, S., and Köster, J. (2021). Sustainable data analysis with Snakemake. F1000Research, 10:33. doi: 10.12688/f1000research.29032.2.

Nica, A. C. and Dermitzakis, E. T. (2013). Expression quantitative trait loci: Present and future. Philosophical Transactions of the Royal Society of London. Series B, Biological Sciences, 368(1620):20120362. doi: 10.1098/rstb.2012.0362.

Obenchain, V., Lawrence, M., Carey, V., Gogarten, S., Shannon, P., and Morgan, M. (2014). VariantAnnotation : A Bioconductor package for exploration and annotation of genetic variants. Bioinformatics, 30(14):2076–2078. doi: 10.1093/bioinformatics/btu168.

Pierce, B. L., Tong, L., Chen, L. S., Rahaman, R., Argos, M., Jasmine, F., Roy, S., Paul-Brutus, R., Westra, H.-J., Franke, L., Esko, T., Zaman, R., Islam, T., Rahman, M., Baron, J. A., Kibriya, M. G., and Ahsan, H. (2014). Mediation Analysis Demonstrates That Trans-eQTLs Are Often Explained by Cis-Mediation: A Genome-Wide Analysis among 1,800 South Asians. PLoS Genetics, 10(12):e1004818. doi: 10.1371/journal.pgen.1004818.

Pritchard, J. K. (2002). The allelic architecture of human disease genes: Common disease-common variant… or not? Human Molecular Genetics, 11(20):2417–2423. doi: 10.1093/hmg/11.20.2417.

R Core Team. R: A Language and Environment for Statistical Computing. R Foundation for Statistical Computing, Vienna, Austria, (2024). URL https://www.R-project.org/.

Schadt, E. E., Monks, S. A., Drake, T. A., Lusis, A. J., Che, N., Colinayo, V., Ruff, T. G., Milligan, S. B., Lamb, J. R., Cavet, G., Linsley, P. S., Mao, M., Stoughton, R. B., and Friend, S. H. (2003). Genetics of gene expression surveyed in maize, mouse and man. Nature, 422(6929):297–302. doi: 10.1038/nature01434.

Simon, M. M., Greenaway, S., White, J. K., Fuchs, H., Gailus-Durner, V., Wells, S., Sorg, T., Wong, K., Bedu, E., Cartwright, E. J., Dacquin, R., Djebali, S., Estabel, J., Graw, J., Ingham, N. J., Jackson, I. J., Lengeling, A., Mandillo, S., Marvel, J., Meziane, H., Preitner, F., Puk, O., Roux, M., Adams, D. J., Atkins, S., Ayadi, A., Becker, L., Blake, A., Brooker, D., Cater, H., Champy, M.-F., Combe, R., Danecek, P., Di Fenza, A., Gates, H., Gerdin, A.-K., Golini, E., Hancock, J. M., Hans, W., Hölter, S. M., Hough, T., Jurdic, P., Keane, T. M., Morgan, H., Müller, W., Neff, F., Nicholson, G., Pasche, B., Roberson, L.-A., Rozman, J., Sanderson, M., Santos, L., Selloum, M., Shannon, C., Southwell, A., Tocchini-Valentini, G. P., Vancollie, V. E., Westerberg, H., Wurst, W., Zi, M., Yalcin, B., Ramirez-Solis, R., Steel, K. P., Mallon, A.-M., De Angelis, M. H., Herault, Y., and Brown, S. D. (2013). A comparative phenotypic and genomic analysis of C57BL/6J and C57BL/6N mouse strains. Genome Biology, 14(7):R82. doi: 10.1186/gb-2013-14-7-r82.

Slatkin, M. (2008). Linkage disequilibrium — understanding the evolutionary past and mapping the medical future. Nature Reviews Genetics, 9(6):477–485. doi: 10.1038/nrg2361.

van der Auwera, G. and O’Connor, B. D. Genomics in the Cloud: Using Docker, GATK, and WDL in Terra. O’Reilly Media, Sebastopol, CA, first edition edition, (2020). ISBN 978-1-4919-7519-0.

Westra, H.-J. and Franke, L. (2014). From genome to function by studying eQTLs. Biochimica et Biophysica Acta (BBA) - Molecular Basis of Disease, 1842(10):1896–1902. doi: 10.1016/j.bbadis.2014.04.024.

White, R. J., Mackay, E., Wilson, S. W., and Busch-Nentwich, E. M. (2022). Allele-specific gene expression can underlie altered transcript abundance in zebrafish mutants. eLife, 11:e72825. doi: 10.7554/eLife.72825.

