## Supplementary Information for "*tadar*: an R/Bioconductor package to reduce eQTL noise in differential expression analysis"

To establish a consensus approach for moderating *p*-values, we evaluated the impact of DAR across five datasets, representing two common model organisms: zebrafish and mouse. A summary of the datasets can be found in Table S1. Dataset 1 is described in Barthelson et al. (2025) and is available in the GEO database (GSE217196). Dataset 2 is described in Barthelson et al. (2020) and is available in the GEO database (GSE151999). Dataset 3 is described in Barthelson et al. (2022) and is available in the GEO database (GSE164466). Dataset 4 is described in Sullivan et al. (1997) and is available at the AD Knowledge Portal (accession number syn20808171, https://adknowledgeportal.synapse.org/). Dataset 5 is available in the GEO database (GSE289151).

In total, 17 comparisons were performed between experimental groups across these datasets. The figures presented here are examples from the comparison between naglu$^{\mathrm{A603fs/+}}$ and wild-type zebrafish, but are considered representative of the overall process. The same procedure was applied to each comparison, and similar trends were observed across all cases. The full analysis is available at: https://doi.org/10.5281/zenodo.14769085.

We began by exploring the distribution of *p*-values across different DAR value ranges. To do so, we divided the list of genes tested for differential expression into quantiles based on their DAR values. We observed that quantiles with higher DAR values contained a larger proportion of genes with significant *p*-values, suggesting that DAR was indeed influencing the classification of genes as differentially expressed (Figure S1). We then hypothesised that in the presence of DAR, *p*-values under the null hypothesis do not follow a uniform distribution, but are skewed towards lower values due to the impact of DAR-associaed eQTLs. Specifically, the observed *p*-value distributions in datasets affected by DAR represent a composite distribution. While a uniform distribution of *p*-values under the null hypothesis is still likely present, it is obscured by the skewed distribution of DAR-biased genes. As a result, the overall distribution more closely resembles a Beta distribution, where the $\alpha$ parameter is less than one, controlling the extent of the right skew. Notably, these DAR-impacted p-values are not drawn from a single Beta distribution, but a composite of Beta distributions where the shape is dependent on the relevant DAR values.

To quantify this, we employed a data-driven approach by performing maximum-likelihood estimation to fit the *p*-values to a beta distribution across various DAR intervals, using the MASS package (Venables and Ripley, 2002). The estimated $\alpha$ parameter from the beta function was found to be highly variable across different DAR intervals, whereas the $\beta$ parameter was consistently estimated at approximately 1. When inspecting regression lines through all fitted values for alpha, a negative linear trend was observed between the $\alpha$ parameter and the median DAR from within each bin for each interval (Figure S2). To further investigate this relationship, we performed linear regression on the $\alpha$ estimates, using median DAR as the predictor variable. Although the trend was consistent across all sets of *p*-values from differential expression testing, the $\alpha$ estimates showed significant variability between datasets, likely due to differing levels of DAR within each dataset and due to different numbers of truly differentially expressed genes within each experiment. To account for this variability, we normalised the $\alpha$ estimates by dividing them by the maximum $\alpha$ estimate within each dataset.

For each dataset, we then used Akaike Information Criterion (AIC) to select the most appropriate model for estimating the normalised $\alpha$ values with DAR, setting the initial model as a quadratic function of DAR. In almost all datasets, the AIC method reported a linear relationship between normalised $\alpha$ and DAR. Linear regression on the normalised $\alpha$ values exhibited a consistent trend across all datasets (Figure S2), where $\alpha$ dropped below $\alpha = 1$ somewhere between DAR values of 0.1 and 0.2. Based on this, we derived a consensus function (Equation 3) to estimate the $\alpha$ parameter using DAR alone, and incorporated a threshold below which $\alpha$ would remain at $\alpha = 1$.

Having devised a model for the $\alpha$ parameter in the context of DAR, we applied this function to directly moderate *p*-values by calculating the cumulative probability of observing the original *p*-value under the modified, skewed distribution. This adjustment is performed independently for each feature tested for differential expression, allowing *p*-values to be moderated to varying degrees depending on the DAR region in which the feature resides. Using this methodology, features only decrease in significance (Figure S3). However, since each feature is moderated independently, the resulting order of the features may differ from the original ranking.

**Table S1. Five datasets were used to assess the impact of DAR and develop an approach for moderating *p*-values.**

| Dataset | Organism | Tissue | Age | Groups |
|---|---|---|---|---|
| 1 | zebrafish | whole larvae | 7 days | psen1$^{Q96K97del/+}$<br>naglu$^{A603fs/+}$<br>wild-type |
| 2 | zebrafish | whole brain | 6 months | sorl1$^{V1482Afs/+}$<br>sorl1$^{R122Pfs/+}$<br>sorl1$^{V1482Afs/R122Pfs}$<br>sorl1$^{+/+}$ |
| 3 | zebrafish | whole brain | 6 months | psen1$^{T428del/+}$<br>psen1$^{W233fs/+}$<br>psen1$^{+/+}$ |
| 4 | mouse | cerebral cortex | 3 months | APOE$^{\varepsilon2/\varepsilon2}$<br>APOE$^{\varepsilon3/\varepsilon3}$<br>APOE$^{\varepsilon4/\varepsilon4}$ |
| 5 | zebrafish | whole brain | 3 months | psen1$^{T428del/+}$<br>psen1$^{W233fs/+}$<br>psen1$^{+/+}$<br>PK/PK<br>Tu/Tu<br>PK/Tu |

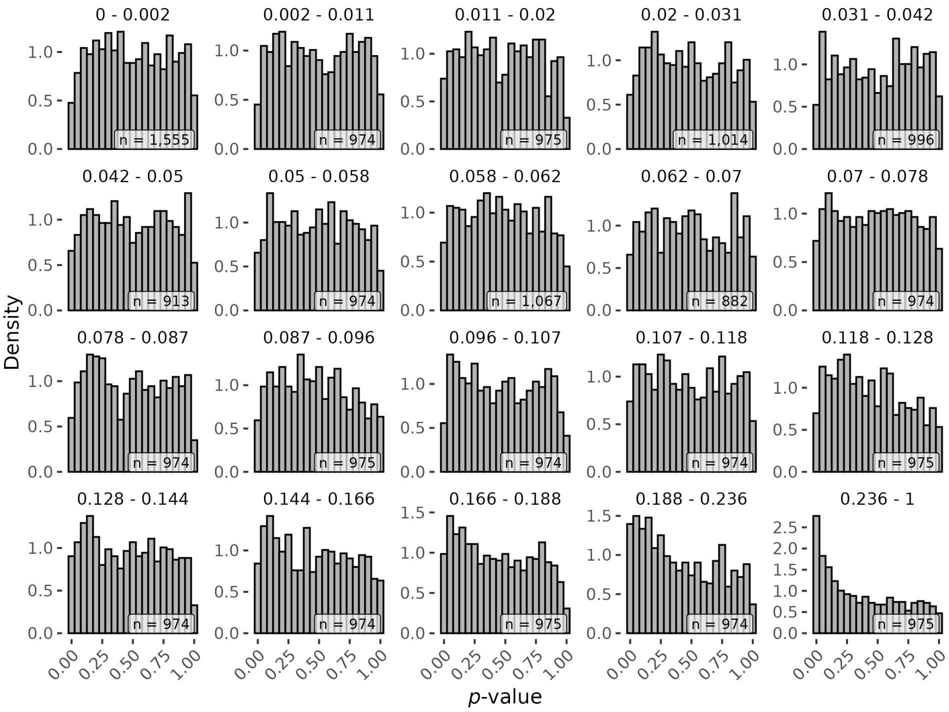


**Figure S1. *p*-value distributions become increasingly right-skewed at higher DAR values.**
Histograms display the distribution of *p*-values from gene-level differential expression testing across 20 approximately equal quantiles established using gene DAR values. The range of DAR values contained within each quantile are indicated above the faceted histograms. The number of genes comprising each quantile are indicated in the lower right-hand corner of each plot.

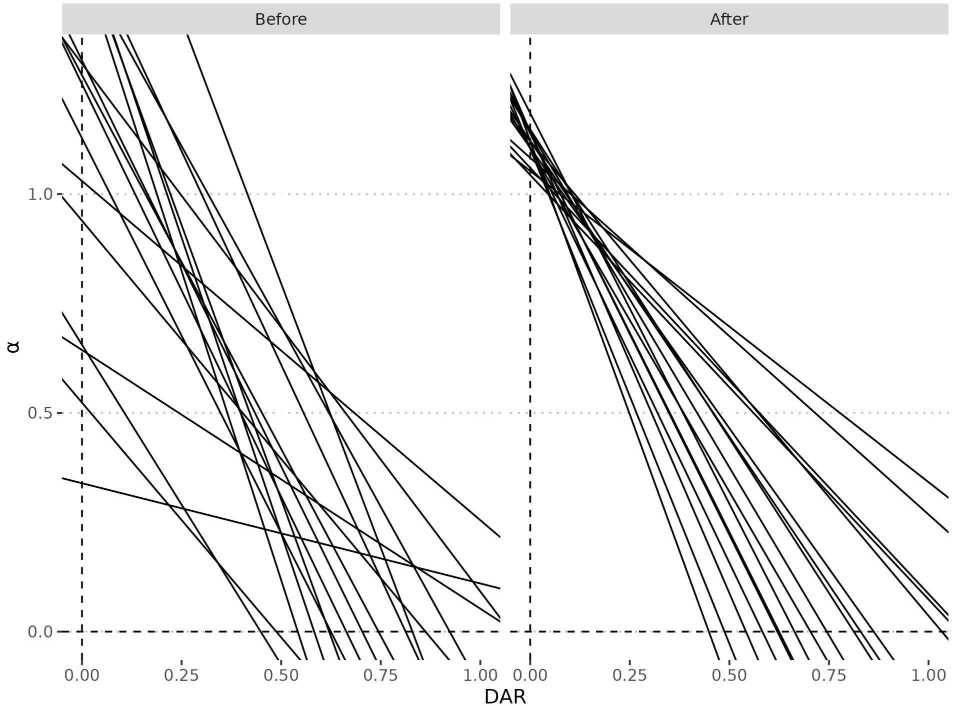


**Figure S2. $\alpha$ as a function of DAR across all 17 comparisons before and after $\alpha$ normalisation.**
Regression lines through fitted values reveal that the relationship between $\alpha$ and DAR for all comparisons across the five datasets follws a linear trend with negative slope. The extent of DAR is dataset-specific, so the estimated $\alpha$ was normalised for each dataset by dividing by its maximum value.

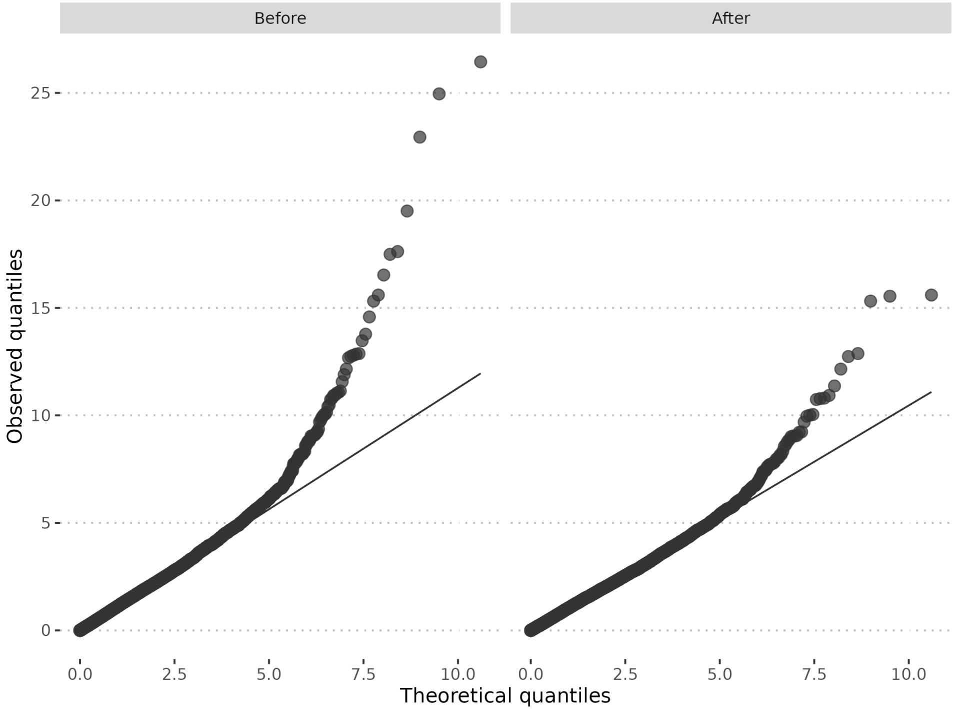


**Figure S3. Quantile-quantile plots before and after DAR modification of *p*-values.**
The negative of the natural logarithm of observed *p*-values were compared against a theoretical exponential distribution to assess how closely they follow the expected uniform distribution under the null hypothesis. Deviation from the $x = y$ relationship indicates rejection of the null hypothesis. Following DAR modification of *p*-values, the observed data more closely resembles the theoretical distribution, displaying an overall reduction in significance across the entire dataset.

### Supplementary References

Barthelson, K., Pederson, S. M., Newman, M., and Lardelli, M. (2020). Brain transcriptome analysis reveals subtle effects on mitochondrial function and iron homeostasis of mutations in the SORL1 gene implicated in early onset familial Alzheimer's disease. *Molecular Brain*, 13(1):142. doi: 10.1186/s13041-020-00681-7.

Barthelson, K., Newman, M., and Lardelli, M. (2022). Brain transcriptomes of zebrafish and mouse Alzheimer's disease knock-in models imply early disrupted energy metabolism. *Disease Models & Mechanisms*, 15(1):dmm049187. doi: 10.1242/dmm.049187.

Barthelson, K., Protzman, R. A., Snel, M. F., Hemsley, K., and Lardelli, M. (2025). Multi-omics analyses of early-onset familial Alzheimer's disease and Sanfilippo syndrome zebrafish models reveal commonalities in disease mechanisms. *Biochimica et Biophysica Acta (BBA) - Molecular Basis of Disease*, 1871(3):167651. doi: 10.1016/j.bbadis.2024.167651.

Sullivan, P. M., Mezdour, H., Aratani, Y., Knouff, C., Najib, J., Reddick, R. L., Quarfordt, S. H., and Maeda, N. (1997). Targeted replacement of the mouse apolipoprotein E gene with the common human APOE3 allele enhances diet-induced hypercholesterolemia and atherosclerosis. *The Journal of Biological Chemistry*, 272(29):17972–17980. doi: 10.1074/jbc.272.29.17972.

Venables, W. N. and Ripley, B. D. *Modern Applied Statistics with S.* Statistics and Computing. Springer, New York, 4th ed edition, (2002). ISBN 978-0-387-95457-8.
